# Diatom community shifts across oligotrophic-eutrophic seasonal gradients and bloom variability as a function of mixing depth in the subtropical Northern Red Sea

**DOI:** 10.1101/2024.04.17.589857

**Authors:** Yoav Avrahami, Gil Koplovitz, Miguel J. Frada

## Abstract

Diatom blooms dominate nutrient-rich ecosystems. Less is known about the ecology and bloom dynamics of diatom populations in oligotrophic ecosystems. Here, we investigated seasonal succession of planktonic diatoms in the Gulf of Aqaba (GoA) at the northern Red Sea. The GoA is a subtropical ecosystem alternating between stratified, oligotrophic profiles during summer, and deeply mixed, mesotrophic during winter. Diatom density and diversity were lower during the stratified season, dominated by pennate species, and increased at mid-winter as nitrate exceeded ∼0.5 µmol L^-1^. Diatom density lagged after total phytoplankton and entailed a transition to centric-diatom dominance, suggesting both higher nutrient requirements for diatom growth and ecophysiological differences between morphotypes. Ephemeral blooms were detected at the mixing-to-stratification transition. Under milder conditions, mixing was shallow and diatoms reached ∼98 individuals. mL^-1^. Small-centric Thalassiosiraceae and several pennates dominated. However, during the following colder year, mixing depth reached ∼700 m. Consequently, nutrient concentrations were higher and diatoms reached ∼390 individuals. mL^-1^. This enabled emergence of chain-forming species (namely *Chaetoceros* and *Leptocylindrus*) along small-centric and pennates, and high spore abundance was detected. Restratification led to rapid bloom decline. These results illustrate diatom community succession and bloom development as a function of nutrient availability in subtropical ecosystems.

## Introduction

Marine phytoplankton form the base of the marine food-web and drive nearly one half of the global primary production (Field *et al*., 1998). Diatoms (Bacillariophyta) are one of the most diverse and prolific phytoplankton groups, responsible for about 40% of the marine primary productivity and acting as a key driver of the biological pump (Nelson *et al*., 1995; Mann and Vanormelingen, 2013; Tréguer *et al*., 2018). Moreover, diatom cell wall (frustule) is primarily composed of silica, thus they also play a primary role in the global silica cycle (Tréguer *et al*., 1995, 2018). Two main morphological groups of diatoms exist based on frustule valve symmetry: centric diatoms with radial symmetry and pennate diatoms that are elongated with primarily bilateral symmetry (Round, 1981; Round *et al*., 1990). Diatom cell-size ranges from a few micrometers up to a few millimeters, where several species form large chains of cells (Fryxell, 1978).

Diatoms are a cosmopolitan group across the oceans (Malviya *et al*., 2016). However, they largely prevail and bloom in well-mixed coastal and upwelling ecosystems characterized by high turbulence and high nutrient loads (Crawford, 1995; Sarthou *et al*., 2005; Ruggiero *et al*., 2018). These blooms form the base of the most important fisheries worldwide (Koeve, 2004; Lassiter *et al*., 2006). As outlined in Leblanc et al. (Leblanc *et al*., 2018), the classical view of diatom spring bloom succession in nutrient rich regions involves a first stage following upwelling or strong mixing that is dominated by smaller fast-growing species (namely of the genera *Thalassiosira* and *Skeletonema*), followed by larger chain-forming species (e.g., of the genus *Chaetoceros*). Then, as the water column becomes stratified and nutrient-limited, species adapted to oligotrophic regimes often carrying N_2_-fixing symbionts thrive (e.g., of the genera *Rhizosolenia* and *Hemiaulus*). This view is possibly skewed to large size species, because the collection of water samples typically relies on large-mesh size plankton nets (∼20-60 um). However, recent studies of diatom blooms report an actual numerical dominance of very small species (< 5 µm; e.g., of the genera *Minidiscus*, *Thalassiosira*, *Fragilariopsis*, *Nitzschia*, and *Nanoneis*), emphasizing their importance during blooms (Savidge *et al*., 1995; Buck *et al*., 2008; Leblanc *et al*., 2018). Diatom ecology and assemblage composition and dynamics in open ocean oligotrophic regions, that constitute in area occupancy the majority of the global ocean, is much less understood. Reports indicate that in oligotrophic regions of the oceans, diatom biomass and diversity is typically low (Nelson *et al*., 1995; Tréguer *et al*., 2018; Busseni *et al*., 2020), although intermittent high-density blooms at the deep- or subsurface chlorophyll maxima (Brzezinski and Nelson, 1995, 1996; Karl *et al*., 2012) and high diversity comparable to eutrophic areas have been reported (Malviya *et al*., 2016; Setta *et al*., 2023).

Here, we examined with high temporal and taxonomic resolution the diversity and succession patterns of diatom assemblages in the subtropical ecosystem of the Gulf of Aqaba/ Eilat (GoA) in the northern Red Sea (Fig. 1). Diatoms were detected by using scanning electron microscopy for detailed quantitative identification. Our overarching goal was to determine with high taxonomic and spatiotemporal resolution the local annual cycle of diatoms and bloom development, including understanding the relative contribution of size-classes and morphological-types (centric vs. pennate species), to foster further understanding on the diversity, patterns of community succession and responses as a function of environmental variability (abiotic) in subtropical ecosystems.

**Figure 1.**
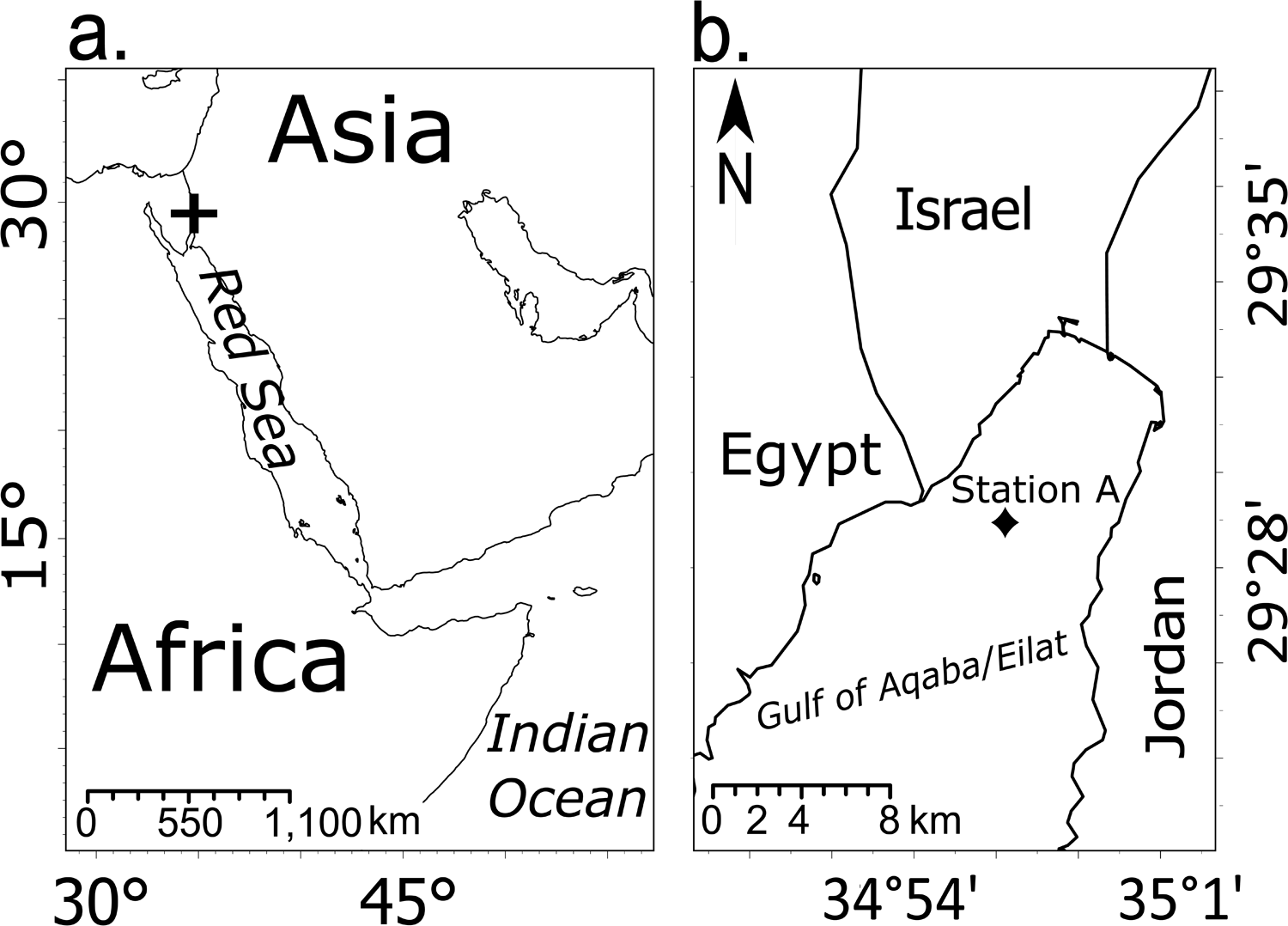
Map of the (a) Red Sea and (b) Gulf of Aqaba (GoA). The Red Sea, connected to the Indian Ocean and the GoA, the northeaster tip of the Red Sea, indicated as plus in a. The ‘Station A’, the open sea sampling station in this study is indicated as filled square in b.

### Study site

The GoA is a warm, narrow basin surrounded by deserts appended to the northern section Red Sea. From an oceanographic standpoint, the GoA features strong seasonal variations in water column stability and consequent nutrient loads. During the warmer spring and summer seasons (April to September) the water column is stratified and oligotrophic, macronutrients are limiting and phytoplankton biomass is low (< 1 µmol L^-1^ nitrate, < 0.15 µmol L^-1^ phosphate, > 0.5 µg L^-1^ chlorophyll a, resembling open ocean gyre ecosystems (Israel National Monitoring Program NMP; Lindell and Post, 1995; Zarubin *et al*., 2017; Keuter *et al*., 2021). Under these conditions, *Prochlorococcus* numerically dominates phytoplankton communities (Lindell and Post, 1995). However, around October surface cooling triggers convective mixing and the erosion of the stratified layer. Continuous cooling through winter drives mixing to a maximum around March. The amplitude of mixing varies largely between years, approximately ranging from about 300m to exceptional depths of >700m during especially cold winters. The depths of mixing vary as an inverse function of atmospheric temperature in winter and is enabled by a prevailing narrow thermocline gradient across the water column (Genin *et al*., 1995; Zarubin *et al*., 2017). Deep mixing transports ample nutrients to the photic layer, resulting in marked shifts in the phytoplankton density and group-composition (e.g., Lindell and Post, 1995; Al-Najjar et al., 2007; Keuter et al., 2023). Then, at the onset of spring during March/April, mixing is capped by surface warming, prompting re-stratification of the water column. This physical state of the water sets the conditions for the rise of a prominent phytoplankton bloom with uniquely high chlorophyll densities for a subtropical ecosystem (Zarubin *et al*., 2017). Large-chain forming diatoms from the genus *Chaetoceros* have been reported to dominate the integrated photoautotrophic biomass during such bloom (Kimor and Golandsky, 1977; Al-Najjar *et al*., 2007). However, comprehensive quantitative assessment of the diatom diversity, succession patterns, bloom development and decline are still lacking. In addition, reports indicate that the GoA is a fast-warming ecosystem (> 0.4°C decade^−1^), far exceeding global averages (0.11°C decade^−1^) (Belkin, 2009; Chaidez et al., 2017).Warming results in shallower winter mixing and substantially weaker blooms (Zarubin *et al*., 2017). Recent studies indicate that local warming selectively impacts coccolithophores, leading to decline of winter species and the emergence of other warm-water, oligotrophic species (Frada *et al*., 2022; Keuter *et al*., 2023). Models have generated contrasting predictions for diatoms, indicate either an increase (Busseni *et al*., 2020) or decrease (Henson *et al*., 2021) due to climate change in oligotrophic open ocean regions. Thus, besides primary goals stated above, we also aimed to generate a reference database for diatoms diversity and ecology in the GoA for future evaluations of the impact of ocean-warming on subtropical ecosystems.

## Methods

### Sampling

Sampling was undertaken during 2020-2022 in the open sea station A (29°28’N, 34°55’E) where bottom depth is about 700 m and is located about 3 km off-shore the city of Eilat (Israel) in the Gulf of Aqaba. During winter mixing (October-March), samples were collected at 2 m depth twice a month. Between March and April, when winter mixing ceased and before full stratification, samples were collected weekly or twice per week to capture bloom events. In the bloom of 2022 sampling took place daily for 5 consecutive days. At each time point, ∼30 L of seawater was sampled using a manual pump installed in a boat and pre-filtered by 500 µm mesh to avoid large zooplankton and transferred into acid washed carboys. In addition, a Sea-Bird SBE 19 CTD (conductivity, temperature, depth, Sea-Bird Scientific) was deployed to obtain physical parameters of the water-column. Mixed layer depth (MLD) was calculated as the depth in which there is a difference of 0.2 □ of potential temperature, compared to 3m depth (Zarubin *et al*., 2017). During summer stratification, samples were collected at the end of April (April 27^th^), May, June, and August, with similar procedures both from the surface but also the DCM using 12 L Niskin bottles (General Oceanics) installed on Sea-Bird carousel and coupled with Sea-Bird SBE 19+ CTD (Sea-Bird Scientific). Within ∼30 min of collection, the seawater samples were sub-sampled for nutrients, pigments, and diatom counts by electron microscopy.

### Nutrient analyses

Total inorganic nitrogen (TIN, NO_2_+NO_3_+NH_4_), silicate (SiO_2_), and orthophosphate (PO_4_) were determined. For the analysis of NO_2_, NO_3_, silicate, and phosphate, sub-samples of 10 mL were transferred in triplicates to 15 mL Falcon tubes and analyzed with Flow Injection autoanalyzer (Quik-Chem 8500, LACHAT Instruments). Every run was calibrated by using commercial standards with at least 5 sub-standards to cover the whole working range. For ammonium (NH_4_) measurements, quadruplicates of 4 mL were transferred to 15 mL Falcon tubes and analyzed fluorometrically (Hoefer).

### Chlorophyll a

Triplicates of 300 mL seawater were filtered onto Whatman glass fiber filters (GF/F, 25 mm diameter, 0.7 µm pore size). The filters were incubated in 90% acetone (Carlo Erba Reagents) buffered with saturated MgCO_3_ (Sigma-Aldrich), over-night (4 □ in darkness), and analyzed fluorometrically for chlorophyll a (Jeffrey and Humphrey, 1975) (Trilogy, Turner Designs).

### Electron microscopy

About 2 L of seawater were gently filtered in duplicates onto polycarbonate filters (Nuclepore Track-Etch membrane, 25 mm diameter, 0.8 µm pore size) and washed with 2mL 0.1mM NaOH to remove salt crystals (possible dissolution of diatom frustules was tested and was undetected). The filters were then dried over-night (room temperature) and coated with a thin Au/Pd film by sputter coater (Cresington 108). Representative sections of each filter were analyzed quantitatively via optical transects at two magnifications by scanning electron microscopy (Phenom Pro): x1,000 for large diatoms (> 50 µm, 100 screens per sample), and x3,000 for small diatoms (< 50 µm, 300 screens per sample). Diatoms were counted as individuals, meaning that a chain containing several cells was considered as one individual (Kenitz *et al*., 2020).

### Data analysis

Diatom community analysis was undertaken by clustering and non-metric multidimensional scaling (NMDS). We subset the data to exclude the effect of rare species and kept only species that met two criteria: a. presence in more than 10% of all sampling points, b. contribution of at least 1% to total relative abundance in at least one sampling point. Other analyses were performed on the entire species list. Species turnover is a temporal analogue measurement of species richness, which intends to highlight the stability of the community over time, by looking on the differences of species composition between two consecutive time-points. This index was undertaken by using the R package ‘codyn’ (Hallett *et al*., 2016), measured as: 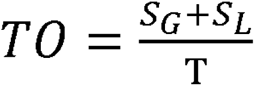. Where TO is total turnover, S_G_ is species gained, S_L_ is species lost, and T is total species observed in both time points. Additional data analyses were undertaken with R software, version 4.2.1 (R Core Team 2021).

## Results

### Temporal variation in physicochemical characteristics of the northern GoA

Diatoms were surveyed from November 2020 to June 2022, encompassing marked temporal variations in physicochemical parameters of the water column (Fig. 2). From November to early April (winter-mixing season), SST progressively declined (Fig. 2a), which drove deep a gradual mixing of the water column over the course of about 5 months. Temperature decline was more important during 2022. As such, maximal Mixed-Layer Depth (MLD) was about 400 m in April 2021 and was > 700 m in April 2022. Right after, SST increased which stopped mixing, setting the stage for water column re-stratification. SST increased from the end of April to September, progressively exacerbating stratification.

**Figure 2.**
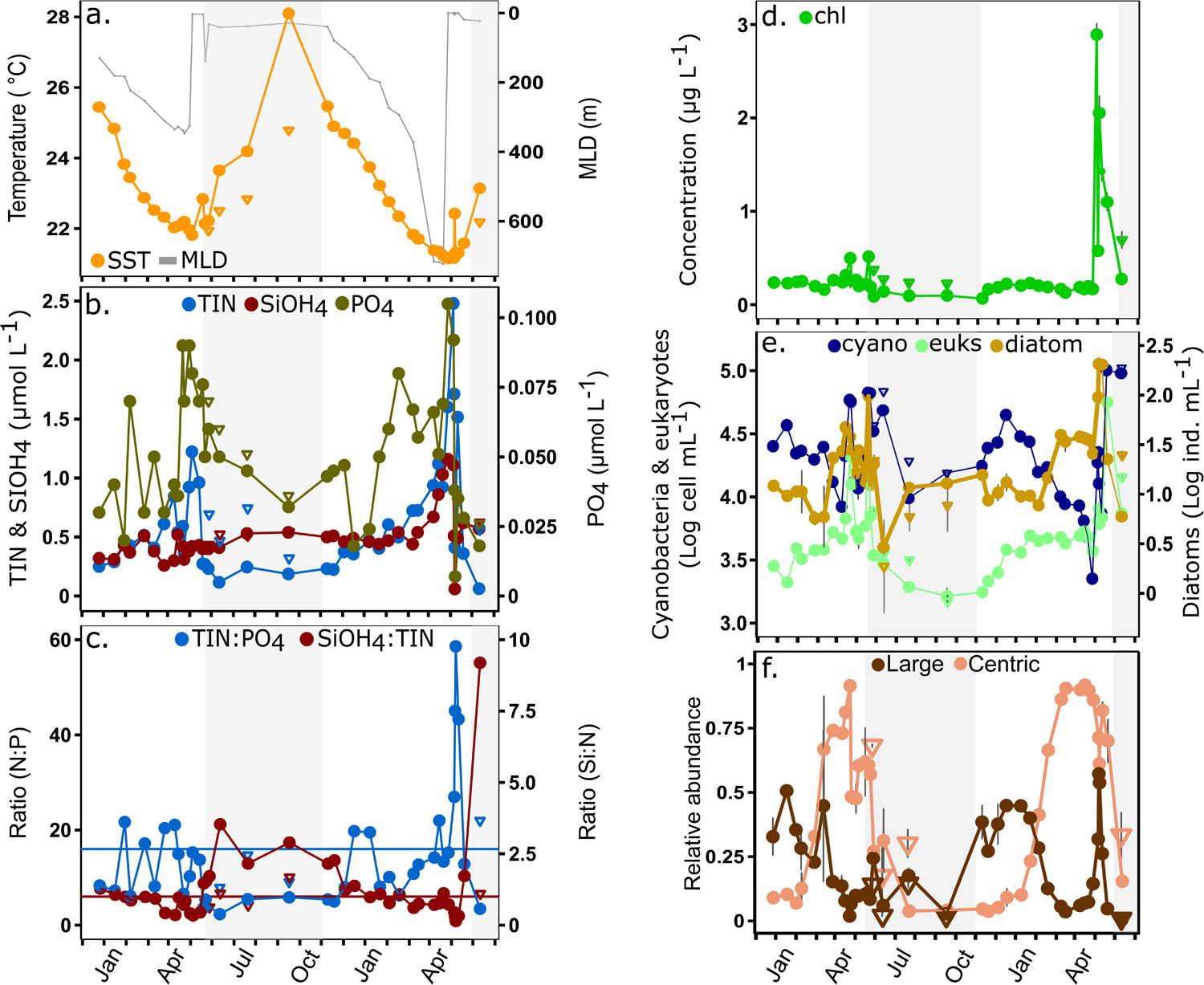
Seasonal dynamics of environmental variables, algal pigments, and algal abundances in the Gulf of Aqaba. The shaded grey area represents the stratified season. Measurements undertaken during the stratified seasons at the DCM are presented in triangle symbols. (a) Temperature (sea surface temperature (SST) and DCM) and the mixed layer depth (MLD). (b) Macronutrients: total inorganic nitrogen (TIN), silicon, and phosphate. (c) Ratios of main macronutrients in the sub-surface: TIN: P and Si: TIN. The horizontal lines represent of Redfield ratio for TIN: P (16:1, blue), and Si: TIN (1:1, red). (d) Chlorophyll a density. Grey bars represent standard deviation (n=3). (e) Averaged log-transformed concentration of total eukaryotic phytoplankton, total cyanobacteria (*Prochlorococcus* and *Synechococcus*) and diatom densities. Grey bars represent standard deviation (n = 2). Data of cyanobacteria and eukaryotes is based on flow cytometry counts, diatoms were counted by scanning electron microscopy. (f) Relative abundance of large diatoms (> 50 µm, compared to smaller diatoms, brown) and of centric diatoms (compared to pennate, pink). Grey bars represent standard deviation (n=2).

The concentrations of the major macronutrients (Total inorganic nitrogen, TIN), orthophosphate (P) and silicate (Si) in surface waters (∼2 m) responded to the seasonal mixing-stratification cycles (Fig. 2b). All macronutrients increased during the mixing season. Maximal nutrient values were concomitant with the MLD peaks in April, particularly in 2022. In 2021, maximal concentration of TIN was 1.2 µmol L^-1^, P was 0.09 µmol L^-1^ and Si was 0.4 µmol L^-1^. In 2022, maximal concentrations of TIN were 2.5 µmol L^-1^, P was 0.11 µmol L^-1^ and Si was 1.2 µmol L^-1^. At the onset of the stratified period, TIN and P declined rapidly to lower concentrations (TIN = 0.2 and P = 0.04 µmol L^-1^ on average). A decline in Si was solely detected in 2022. During the stratified seasons, higher TIN and P were detected at the DCM as compared to the surface. The concentrations of Si during summer were comparable or higher than in winter 2021, but lower than in winter 2022.

The ratios between macronutrients highlight variations in their relative availability to phytoplankton growth (Fig. 2c). The ratio of TIN:P of 16:1 (Redfield *et al*., 1963) and Si:TIN of 1:1 (Brzezinski, 1985) were used as reference levels. Rapid fluctuations in TIN:P ratio between 5.8 and 22 were detected during the mixing seasons. Peak TIN:P ratios of ∼60 was detected in April 2022. During the stratified seasons, TIN:P was low averaging 4.2 t the surface. By contrast, Si:TIN was always 3.9 on average during stratified seasons (Si: TIN), but it progressively declined below 1:1 during the mixing season with minima in April (0.15 on April 7^th^ 2022). Si: TIN bounced back to higher values between ∼3 and >9 in May 2021 and 2022, respectively.

### Phytoplankton and diatoms

Chlorophyll *a* (Chla) concentration varied in direct response to the mixing-stratified seasonal cycles as well (Fig. 2d). Peak Chla concentrations were detected in April, particularly in 2022 (2.89 µg L^-1^) as compared to 2021 (0.51 µg L^-1^). We defined this peak in Chla during April as the bloom. Chla during the stratified seasons was on average 0.14 µg L^-1^ at the surface and 0.36 µg L^-1^ at the DCM.

Total photoautotrophic cells (eukaryotes and cyanobacteria) were determined by flow cytometry (Fig 2e, Suppl. Fig. 1). Maximal photosynthetic eukaryotes densities broadly coincided with the chlorophyll peaks. About 27.5 × 10^4^ cells mL^-1^ were detected in April 2021 and 56.1 × 10^4^ cells mL^-1^ in April 2022. During the stratified seasons eukaryotic concentration were a magnitude lower, averaging 3.5 × 10^3^ cells mL^-1^. A rapid increase in cell concentrations were detected during the onset of the mixing season. Cyanobacteria concentrations declined during the mixing seasons and peaked right after the eukaryotic bloom in April/May.

Diatoms were quantitatively identified by SEM. Single cells or cell-chains were both counted as single individual (Fig. 2e). Diatom average concentrations (individuals mL^-1^, Suppl. Fig. 2) was ∼10 individuals mL^-1^ during the stratified season at the sea surface were and ∼13 individuals mL^-1^ at the DCM. During the winter-mixing period until about January, diatom concentrations remained comparable to summer. This represented a response-delay relative total Chla and total eukaryotic phytoplankton concentrations (see above). However, around January and February diatom density increased to about 20 individuals mL^-1^. This coincided in both years with MLD of about 300m and an increment in TIN concentrations above 0.5 µmol L^-1^. Maximal densities in diatoms were detected in April, coinciding in both years with the peak in nutrient and chlorophyll concentrations (bloom). Diatom peak concentration during the bloom was four-times higher in April 2022 (393.8 ± 50.7 ind. mL^-1^) relative to April 2021 (98.4 ± 11.2 ind. mL^-1^). After, diatom maxima declined rapidly to minimal concentrations in May (12 and 6 ind. mL^-1^ in 2021 and 2022, respectively). Overall, by comparison to total eukaryotes, diatoms constituted less than 1% of cell counts in most yearly time points, with the exception of the spring bloom during 2022 where diatoms reached ∼6% of eukaryotic cell counts (Suppl. Fig. 3).

Broad size-class analyses (Fig. 2f) indicated clear seasonal variations in the abundance of larger diatoms > 50 µm (which include chain-forming individuals) and smaller diatoms < 50 µm. During summer 80-95% of diatom were < 50 µm in size. At the onset of mixing about size categories shared about 50% of the assemblages from October to January. From January to the bloom, larger diatoms declined and smaller diatoms comprised >90% of the diatoms. During the blooms larger diatoms increased again. In April 2021 larger cells reached up to 25% of diatoms. In April 2022 larger cells reached up to 60% of diatoms. The relative contribution of centric and pennate diatoms also varied seasonally (Fig. 2f). Pennate cells dominated during summer (>95%). Centric diatoms increased between January and bloom periods, comprising up to 90% of diatoms counts during March and April. A rapid decline in centric species was detected after the blooms, namely following the peak bloom in April 2022.

### Diatom diversity, richness and turnover

SEM analyses enabled genus or species-level taxonomic resolution for most centric diatoms, with the exception of small centric species from the family Thalassiosiraceae. In contrast, most pennate diatoms, particularly with lightly silicified frustules, could not be taxonomically identified by SEM, with the exception of *Cylindrotheca* spp. and *Pseudo-nitzschia* spp. Henceforth, for quantitative purposes all other pennate diatoms were unified in three broad size sub-categories (Large: > 50 µm; Medium: 20-50 µm; and Small: < 20µm). Representative SEM imagery of the main species is presented in Fig. 3.

**Figure 3.**
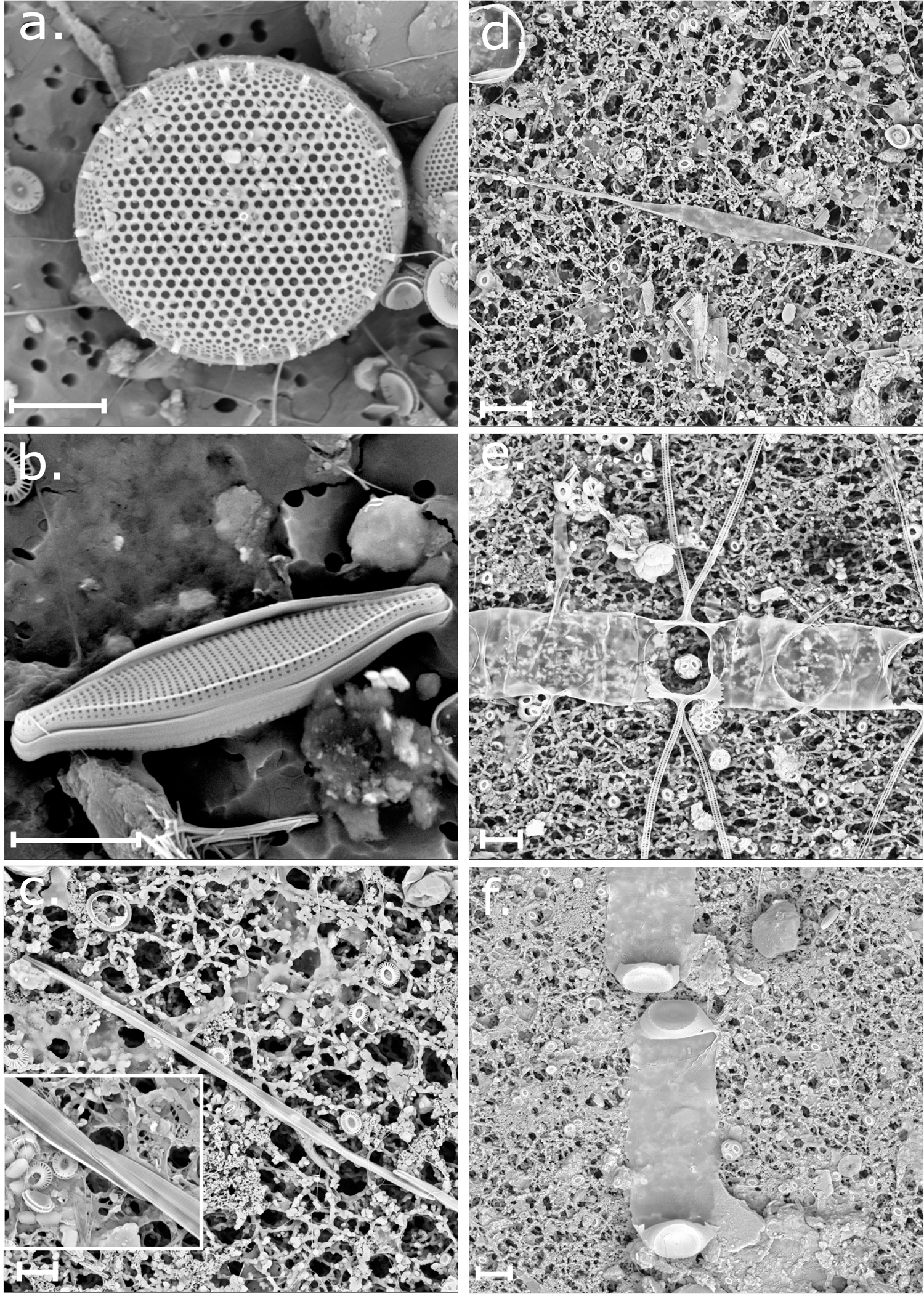
Main diatom species in the GoA. (a) centric diatom from Thalassiosiraceae family; key contributor to the small size-fraction. (b) An unidentified pennate. This group was classified into 3 size-based groups (both in the small and large size-fractions). (c) Chain of the pennate *Pseudo-nitzschia* spp., attachment of valves is shown in inset. (d) A pennate *Cylindrotheca* spp., which appeared in both small and large size fractions. (e) Chain-forming centric *Chaetoceros*; notable representative of the large size-fraction during the bloom. (f) A chain-forming centric *Leptocylindrus*. Scale = 5 µm (a-c), 10 µm (d-f).

Overall, *Cylindrotheca* spp. and other pennate cells were the most common diatoms during summer and early-mixing period (Fig. 4). Pennates decline in the mid-mixing period (around January), concomitant with increase of Thalassiosiraceae. Peak concentrations of Thalassiosiraceae were detected during blooms, particularly in 2022 where they reached up to 145 individuals mL^-1^. Large chain-forming centric species *Chaetoceros* spp. and *Leptocylindrus* spp. were detected in most samples at low concentrations (< 3 individuals mL^-1^). However, during the bloom of April 2022 peak concentrations of 39.5 and 83.7 individuals mL^-1^, respectively, were detected. A few other chain forming species were also detected in April 2022 such as *Rhizosolenia* spp., *Helicotheca sinensis*, *Lauderia annulata* and *Skeletonema grevillea* (Fig. 4). The pennate *Pseudo-nitzchia* spp. was overall rarer, but a clear rise in concentration was also specifically detected during April 2022 (Fig. 4).

**Figure 4.**
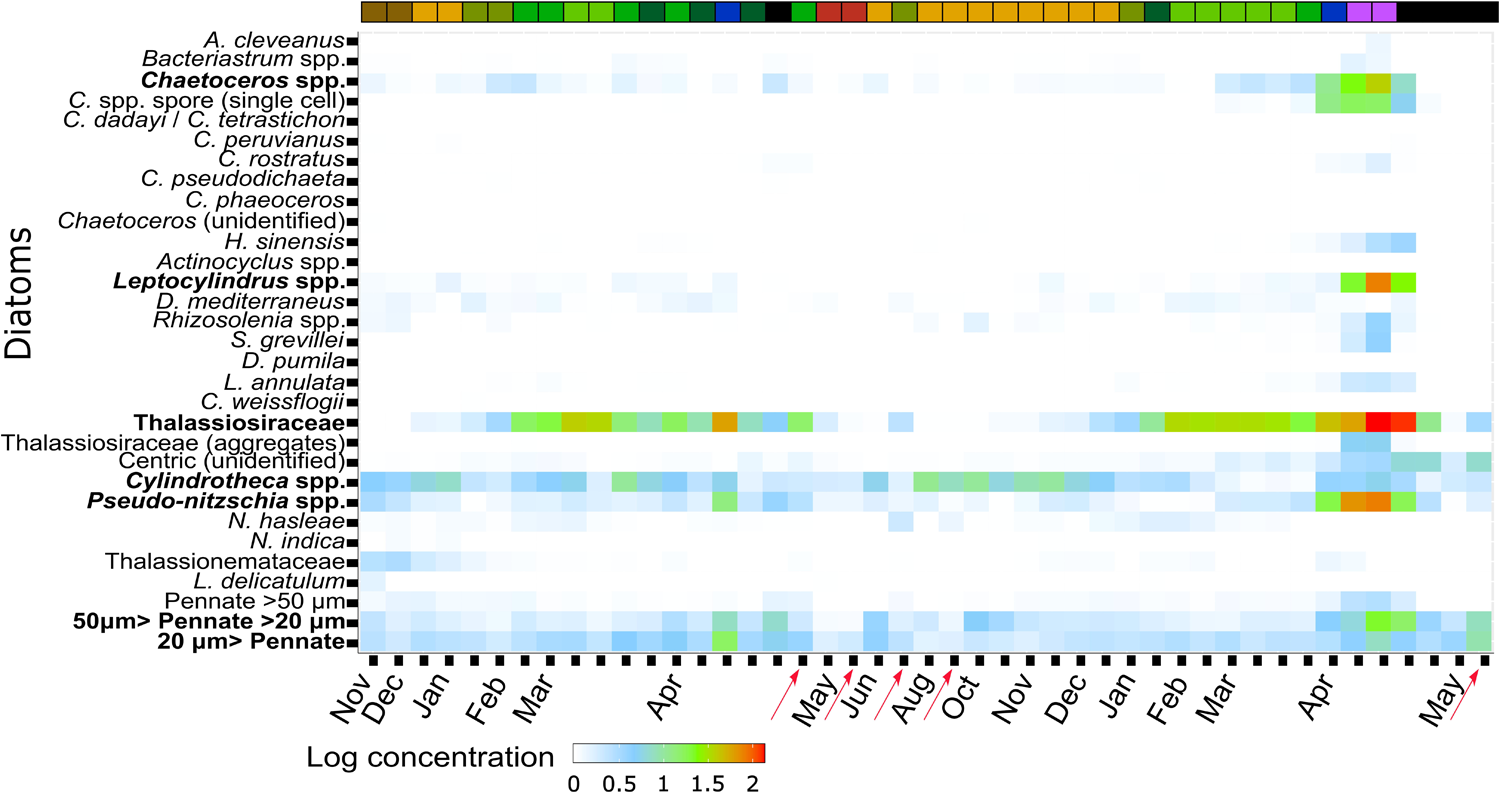
Heatmap of diatom groups composition between 2020-2022. White-red gradient represents log-transformed concentrations (electron microscopy counts). The red arrows represent DCM samples. The color code on top of the heatmap refers to the diatom community cluster, based on 65% similarity (Bray-Curtis index. For further information see Suppl. Fig. 5).

Bray-Curtis similarity analysis was used to outline the main diatom communities. Nine clusters of multiple samples were identified at a 65% similarity level, while five samples were non-clustered (Suppl. Fig. 4). The main species in each cluster or non-clustered sample are specified in Suppl. Table 1. NMDS analyses enabled to link these communities to main oceanographic variables (Fig. 5). Color coding for each community cluster is present in Fig. 4. Broadly, communities X1 and X2 occupied the summer and early mixing periods associated with higher temperatures and low nutrients. These were similar and comprised namely *Cylindrotheca* spp., pennates smaller than 50 µm, and to lesser extent *Pseudo-nitschia.* spp., and. Then communities X3 to X6 occupied most of the period from January to March as nutrient loads increased. Here, Thalassiosiraceae and various pennate species were the most important diatoms. During the bloom Thalassiosiraceae and pennates prevailed in 2021. However, in 2022 *Chaetoceros* was also important. Bloom samples associated with high TIN concentrations and high TIN:P ratios. The post-bloom samples collected in May 2022 associated with low nutrients, but high Si:TIN ratio, included community X9 and several non-clustered samples, where *Cylindrotheca* spp. was the most common diatom. The samples collected in the DCM during the stratified periods were scattered in different clusters. During the early stratified periods, DCM samples were compositionally closer to late winter samples (X3, X5), while in August DCM communities resembled more surface samples from the same period (X1) and the early-mixing period (X2).

**Figure 5.**
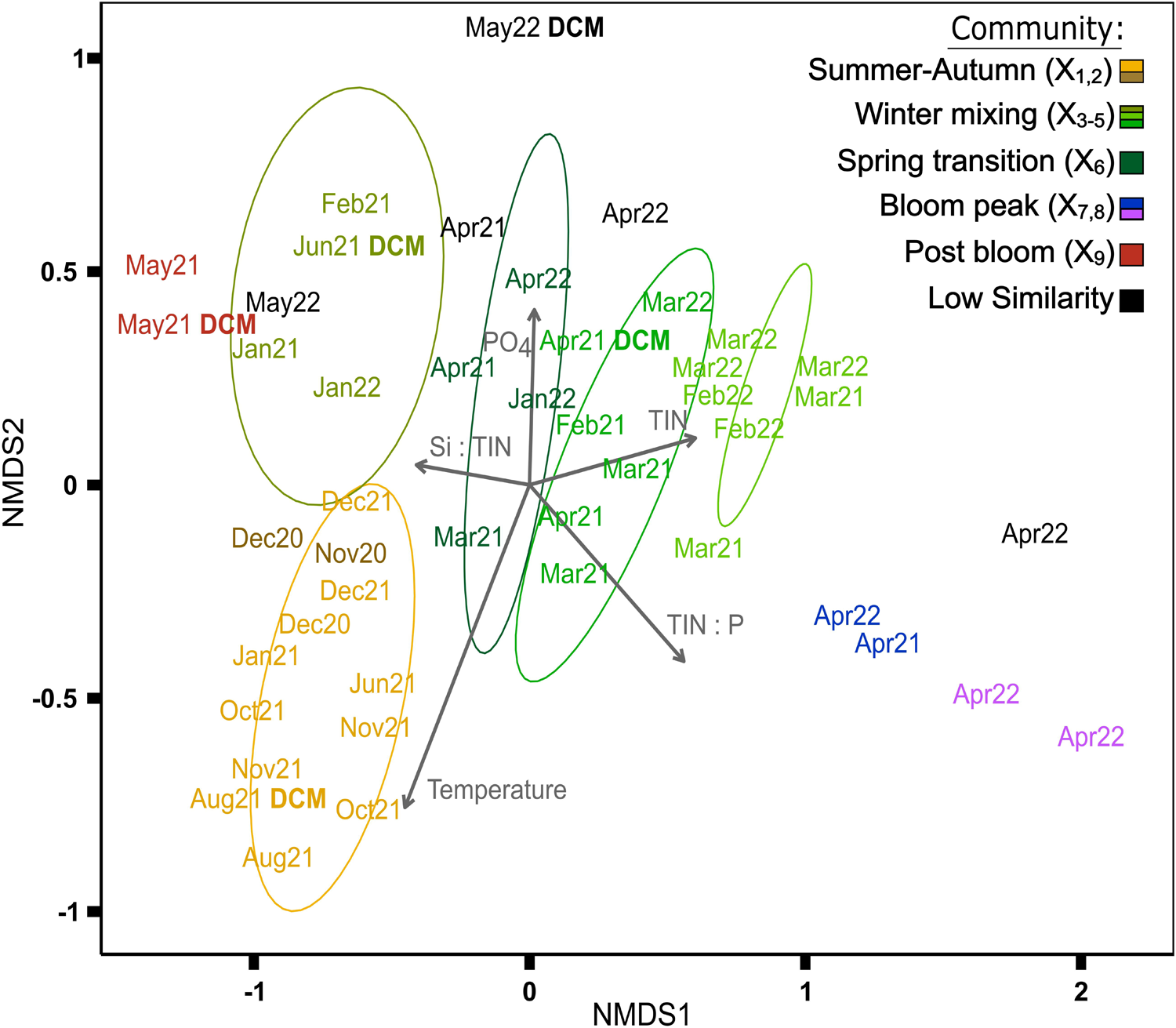
NMDS ordination of diatom communities (Bray-Curtis distance). The color code shows communities with ≥ 65% similarity and represent clusters: Orange, brown: summer and autumn. Light green (3 sub-clusters): winter mixing. Dark green: transition from mixing to stratification. Blue, pink: spring bloom. Red: post bloom. Black shows communities with < 65% similarity to other samples. Grey arrows represent significant environmental variables (p < 0.05).

Diversity indices, Richness and Shannon (H’) were determined of the taxonomic resolution and size-class groupings obtained (Fig. 6a). Broadly, Richness and H’ were higher during the mixing and bloom periods (Richness >15; H’ >1.5), and rapidly declined after the bloom to minimal values in May (Richness <10; H’ < 1.5). Finally, turnover analysis was undertaken to identify period of rapid fluctuation in diatom diversity (Fig. 6b). To circumvent the overrepresentation of samples during winter, a representative sample per month was used for the analysis. The highest turnover was detected after the blooms, particularly in 2022 (∼0.9). The other time points oscillated around 0.3. Minimal turnover was detected in February 2022 at ∼0.1.

**Figure 6.**
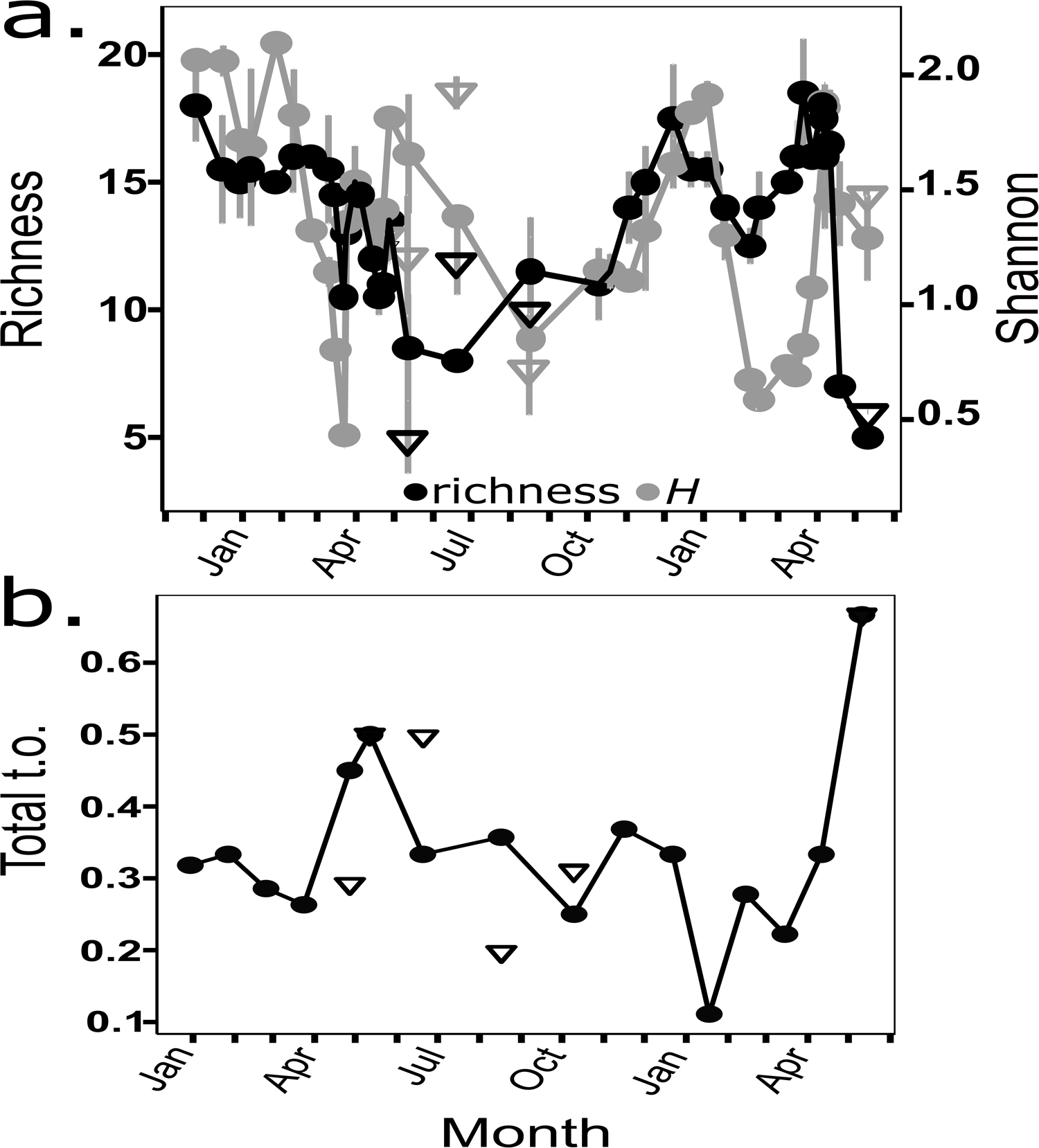
Diatom diversity indices. Circles connected with line represent 2m depth, while triangles represent samples from the deep chlorophyll maximum (DCM). (a) Diatom species richness (black) and Shannon index (grey). Grey bars represent standard deviation (n=2). (b) Diatom species turnover index. This index detects differences between time points in appearance and disappearance of diatom taxa.

### Spore detection and distribution

Spores from members of the chain forming *Chaetoceros* spp. were detected during the mixing and bloom periods (Fig. 7). Interannual variations in spore density were large. In 2021 spores were rare. A few spores were detected in February when diatom concentration increased during mixing, and a few more during the bloom peak in April (∼3-9% of total *Chaetoceros* spp.) (Fig. 7f). In 2022 spore density was much higher. Spores were detected in March during the late mixing period (∼4-17% of total *Chaetoceros* spp.) and then increased even further during the bloom to about 17 spores per mL, accounting for 30-50% to the density of vegetative *Chaetoceros* spp. cells (Fig, 7g). Spores were mainly detected isolated, but also within the cell chains during the bloom in 2022. In April 19th 2022, after the bloom declined, the density of spores was much lower, but vegetative cells were completely absent and thus spores comprised 100% of *Chaetoceros* spp. individuals.

**Figure 7.**
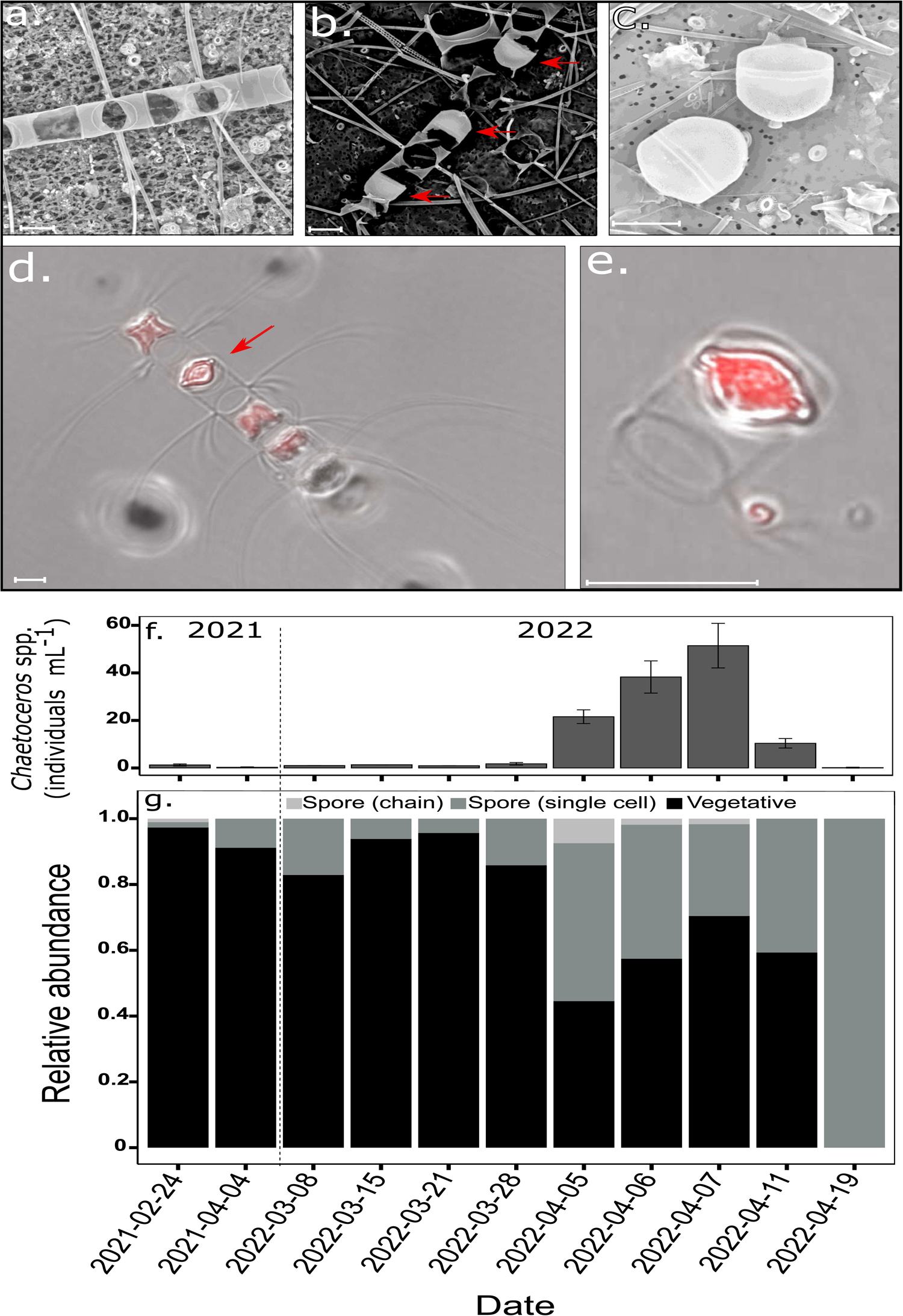
Life-stages of *Chaetoceros* spp. via scanning electron microscopy (a-c) and light-microscopy (d,e). (a) Chain of vegetative cells. (b) A chain, including spores (red arrows) and a vegetative cell. (c) Spores as single cells. (d) A chain, including a spore (red arrow) and vegetative cells. (e) Spore as a single cell. Two layers are presented in light-microscopy: Brightfield (grey) for cell morphology, and chlorophyll autofluorescence (red, EX:649 nm/EM:670 nm). Scale: 10 µm. (f) Concentration of total *Chaetoceros* spp. individuals (including vegetative and spore life forms) by SEM counts. Bars represent standard deviation (n=2). (g) Relative abundance (between 0-1) of vegetative cells and spores of the chain forming *Chaetoceros* spp. species complex, in samples where spores were detected. Black= vegetative cells, grey= spores found as a single cell, light-grey= spores found in a chain.

## Discussion

Here we examined by scanning electron microscopy the diversity and seasonality of diatoms in the GoA with emphasis on bloom development in order to: 1) expand descriptions on diatom diversity and community succession as a function of environmental variables in subtropical ecosystems and, 2) generate a baseline information for diatoms in an ecosystem undergoing rapid warming. In the following paragraphs, we discuss the main ecological patterns observed.

Marked shifts in diatom density and diversity were detected through the annual cycle in the GoA. These shifts were namely associated with seasonal transitions between the stratified, oligotrophic periods and the winter-mixing season. Diatom density was higher during the winter-mixing and specially bloom periods associating with higher nutrient availability and lower temperature and were much lower during the stratified periods characterized by warmer conditions but lower nutrient availability. Diatom species richness and diversity followed parallel patterns. Given limitation in identifying all diatoms detected, especially pennate cells that were largely overlooked due to difficulties in determining precise taxonomic affiliations, species richness and diversity indices were lower during the oligotrophic seasons relative to the nutrient-rich, winter-mixing and bloom periods. This diversity variations between seasons follow analogous patterns described for diatoms in other regions in microscopy-based time-series studies (Hulburt *et al*., 1960) as well as some DNA barcoding-based offshore-near-shore transect gradients (James *et al*., 2022) and large-scale assessments if diatom diversity patterns were richness associated with eutrophic regions (Busseni et al. 2020; but see Malviyia et al. 2016 and Setta et al. 2023 for contrasting patterns). However, it contrasts with general patterns of phytoplankton diversity in the ocean that typically increase toward oligotrophic regions (Righetti *et al*., 2019) and local diversity patterns observed for other phytoplankton as coccolithophores (Keuter *et al*., 2023) This might be explained by overall distinct ecophysiological attributes of diatoms, as r-strategist and selected to thrive in nutrient-rich, turbulent regimes as compared to other phytoplankton groups (Margalef, 1978; Kemp and Villareal, 2018). Yet, as we see in the GoA they also thrive in oligotrophic conditions, without clear signs on nutrient inputs that could sustain diatom standing-stocks. We believe that although pennate diversity was poorly resolved, the same overall pennate-morphotypes were detected also during the winter-mixing. Thus, the diatom richness and diversity patterns shown here are highly plausible. Future DNA-barcoding approach will enable to solve uncertainties. In concomitant, higher efforts to isolate and generate reference datasets in particularly for much less represented pennate species, would largely improve the resolution of taxonomic identification and diversity estimation with optical and molecular methods available.

We note that relatively small pennate diatoms were the most common cells during the stratified period in the GoA, while centric diatom were numerically rarer. Specifically, both *Cylindrotheca* and *Pseudo-nitzschia* were the most common pennate cells. In contrast, centric diatoms rose to numerical prevalence during deep-mixing periods in mid-winter and the bloom. This suggest both that pennate diatoms may be better apt to ensure warm, oligotrophic periods and centric diatoms require higher nutrient, turbulent and possible colder conditions to flourish. Due to high surface-area-to-volume ratios, nutrient uptake kinetics in small cells are advantageous in low nutrient settings relative to large cells (Key *et al*., 2010). This advantage may be even further incremented in pennate cells with fusiform shapes and potentially better apt to compete under limiting conditions (Leynaert et al., 2004; Timmermans et al., 2004; Karp-Boss and Boss, 2016), particularly at high Si:N ratios (Sommer, 1998) as we detected during summer in the GoA. This can be reciprocally supported by higher presence of centric cells in the GoA during April and June at the DCM, where nutrient concentrations are higher, relative to the surface. Noticeably, pennate diatoms are typically viewed as predominantly benthic, colonizing intertidal and well-lit subtidal sediments (Underwood and Kromkamp, 1999). However, as we see in the GoA and reported in other studies, they can be prevalent in planktonic assemblages in oligotrophic settings (e.g., Hulburt et al., 1960, Setta et al. 2023), which may be explained by nutritional advantages stated above, but possibly also by heterotrophic capabilities as it has been indicated for *Cylindrotheca* (Lewin *et al*., 1970), symbiosis with nitrogen-fixing bacteria (Schvarcz *et al*., 2022; Tschitschko *et al*., 2024), or possibly higher thermal maxima. Potential ecophysiological differentiation between diatom morphological types and the exploration of inherent physiological and genetic attributes enabling adaption to oligotrophic conditions in pennate diatoms relative to centric diatoms need further mechanistic investigation.

An additional observation depicted in our study was the fact while total phytoplankton (Chlorophyll a, and eukaryotic phytoplankton cells detected by flow cytometry) rapidly increased within less a month to mixing and initial increased fluxes of nutrients to the photic layer, diatom density and community structure remained relatively unchanged until mid-winter (January). Diatoms are typically capable of responding positively to nutrient inputs and to rapidly overtake phytoplankton communities (Margalef, 1978; McNair *et al*., 2018). Thus, a rapid response of diatoms to early mixing would be expected. We note in addition that the increase in diatom concentration in mid-winter coincided with a marked shift in community composition. This involved in one hand a marked increase in the concentration of small centric Thalassiosiraceae. Incidentally, this coincided with periods when mixing deepened below 300 m as a result of temperature decline below ∼24 □, and TIN concentration exceeded 0.5 µmol L^-1^, suggesting a key selective role on nutrients. Centric diatom cells have been shown to display high half-saturation constant for nutrients (Eppley and Thomas, 1969; Litchman *et al*., 2007) including relative to cohabiting pennate diatoms cells of similar size (Leynaert *et al*., 2004; Timmermans *et al*., 2004). Thus, it is plausible that diatoms, specifically Thalassiosiraceae, experience nutrient limitation during early mixing relative to co-occurring phytoplankton, resulting in a delayed response of diatoms to the winter-mixing dynamics. Lower temperature levels may be a key determinant factor enabling growth of Thalassiosiraceae later in winter. The influence of both factors on centric diatoms ought to be investigated. On the other hand, reciprocal decline of the pennate *Cylindrotheca* was also detected over the same period. This decline was however temporary since *Cylindrotheca* increased again during the bloom at the onset of stratification, although seawater temperature was lower and nutrient concertation were higher. Recent work in the GoA indicates that phytoplankton communities experience light limitation in the GoA as a consequence of deep convective mixing during winter inhibiting growth (Berman and Gildor, 2022; Berman *et al*., 2023). Thus, it is plausible that *Cylindrotheca* is inhibited during deep-mixing periods, while Thalassiosiraceae with eventually higher light absorption efficiencies is unaffected by extended dark periods during deep convective mixing.

Peak diatom densities in the GoA were detected in April during the mixing-to-stratification transitional period as part of the total phytoplankton bloom (total chlorophyll a peak). This agrees with previous detailed reports, indicating that once mixing stops, phytoplankton cells are no longer vertically dispersed through the deep mixed layer and accumulate in the upper surface layers with less nutrient and light restrictions to form a spring bloom (Zarubin *et al*., 2017; Berman *et al*., 2023). A much more prominent bloom was detected in 2022 in direct relation to a much deeper MLD and ampler nutrient fluxes to the surface (Zarubin *et al*., 2017). Particularly small centric Thalassiosiraceae, but also *Pseudo-nitzschia* and *Cylindrotheca* as well as other pennates were the most frequent diatoms during blooms, indicating the importance of small-size species for diatom production during blooms (Leblanc *et al*., 2018). Chain-forming cells were rare in the shallow bloom of 2021. However, concomitant to deeper mixing and ampler nutrients, large chain forming species were also distinctly common in the 2022 bloom, namely *Chaetoceros and Leptopcylindrus*. This observation, links large chain-forming species to nutrients, suggesting lower nutrient availability during shallow mixing limits the growth of large diatoms with higher nutrient requirements, while deep mixing enables bloom development. In addition, light availability may also play an important role in restricting large chain-diatom blooms to the mixing-stratification transition period. Previous photophysiological studies have shown that large diatom cells have lower pigment-specific absorption efficiencies relative to smaller diatoms (Finkel, 2001; Finkel *et al*., 2004). Thus, light limitation during deep convective mixing may inhibit large diatom cells.

Noticeably, diatom blooms declined markedly towards the month of May as stratification was established. This corresponded to the strongest decline in diatom diversity and highest species turnover rates detected in our survey (particularly 2022). Large-chain diatoms were mostly removed and smaller cells declined to low densities. The bloom peak coincided with Si:TIN minima and the decline to large depletion of all macronutrient pools, namely TIN, suggesting a prevalent role of nutrient limitation driving diatom bloom collapse. This assessment agrees with recent integrated model of phytoplankton blooms in the GoA (Berman and Gildor, 2022). This can be further facilitated by rapid export of large chain-diatoms and heavy diatom aggregates at the onset of stratification (Alldredge and Gotschalk, 1989; Grimm *et al*., 1996; Kemp *et al*., 2000), explaining the more prevalent decline of larger cells. Biotic factors such as grazing and viruses are often determinant to diatom decline in high-nutrient regions(Krause *et al*., 2010; Kranzler *et al*., 2019, 2021). These can have a selective effect of communities and underlie the decline of dominant species. In our study we did not assess the contribution of biotic factors. However, these require further investigation to determine the relative prevalence of bottom-up and top-down controls of diatom blooms in the GoA.

As part of our observations, diatom spores associated to the chain forming *Chaetoceros* group were detected during the winter mixing and bloom in both years studied. Spores were also much more common in 2022. This is the first report of diatom spores in the Red Sea. Most of the detected spores appeared isolated but also within cell-chains, indicating recent formation. Diatom spores are often produced in response to adverse environmental conditions for vegetative growth, namely nitrogen limitation (Sugie and Kuma, 2008), decline in light intensity (Smayda and Mitchell-Innes, 1974; Doucette and Fryxell, 1983), fluctuations in salinity (Oku and Kamatani, 1997) and/or temperature (McQuoid and Hobson, 1995) and can serve as seedbank to subsist in a dormancy or quiescent state under sub-optimal (Smetacek, 1985; Ellegaard and Ribeiro, 2018). The triggers of spores for diatoms in the GoA are unknown. However, it may be that nutrient limitation and light fields that vary considerably across the mixing and bloom seasons play a determinant role. Particularly, TIN: P levels were often below the Redfield ratio through winter and a progressive decline in the Si:TIN levels also below the Redfield ratio was also detected, indicating some degree of nutrient limitation during the winter season. Additionally, density-dependent cues particularly viral infections have been also linked to sporulation in *Chaetoceros* (Pelusi *et al*., 2020) and could have played a key role in particularly in 2022 as diatom concentration was much higher and potential host-virus interplay more frequent. Given the importance of spores in diatom ecology, the key determinants for diatom sporulation, their importance for survival as well as contribution to diatom standing-stocks in the water column warrant further examination.

## Conclusions

The GoA represents to some extent a hybrid ecosystem defined by marked seasonal transitions between an oligotrophic and eutrophic regime resembling oceanic gyre and upwelling ecosystems supporting phytoplankton blooms, respectively (Lindell and Post, 1995; Zarubin *et al*., 2017). This oceanographic context offers the opportunity to study subtropical phytoplankton communities but also their responses to environmental changes across seasonal gradients. In our study, we detected clear shifts in diatom communities’ diversity and density through the annual cycle in the GoA. These were namely associated with the development of mixing and entrainment of nutrients during winter and sharp restratification during spring. Overall, we found that small cells are numerically prevalent across seasons, supporting recent reports on the prevalence and importance of small diatoms overlooked in classical studies (e.g., Leblanc et al., 2018; Bolaños et al., 2020; Setta et al., 2023). Pennate species composed most of the summer diatom communities, while centric diatom cells only gain prevalence in later winter and bloom as nutrients increased and temperatures decreased. This points towards marked ecophysiological differentiation between diatom morphological types that need further exploration. We also found that nutrient availability set by the depth of the annual mixing determines both magnitude and community structure of diatom blooms. Specifically, the prevalence of large cells depends on deep-mixing and consequent high nutrient flux to the photic layer, which may be plausibly explained by higher nutrient requirements for growth of large cells relative to smaller cells. These interannual variations in community structure and predominance of large cells can have important cascading consequences enhancing nutrient uptake and carbon export dynamics in marine ecosystems (Bergkvist *et al*., 2018; Tréguer *et al*., 2018). Thus, they can have critical implications for local biogeochemistry and higher trophic level production that warrant further examination. Diatom blooms detected in the GoA are yet of much lower magnitude than reports from high latitude and coastal areas (Koeve, 2004), but follow similar boom-bust dynamics with rapid rise and decline (Brzezinski *et al*., 2001), suggesting tight bottom-up and top-down controls. We also detected recurrent spore production during the mixing and bloom periods, clearly indicating the importance of spores for survival and ecological dynamics of diatoms in subtropical regions. Spore formation can also have additional impact in mass export events, due to heavily silicified frustules that rapidly sink and further enhancing the biogeochemical impact of large diatom cells (e.g., Salter et al., 2012; e.g., Rynearson et al., 2013). To conclude, our study provides a detailed overview of diatom diversity and ecological community succession in a model subtropical ecosystem with high temporal and taxonomic resolution. Given the importance of subtropical regions and their projected expansion in coming decades due to global warming, understanding the patterns of phytoplankton diversity and responses to environmental variations is key to forecast phytoplankton community composition and resilience in the future oceans.

## Acknowledgements

We thank the Israel National Monitoring Program at the Gulf of Eilat for sharing and discussing environmental data. We thank Diana Sarno and Wiebe Kooistra from Stazione Zoologica Anton Dohrn Napoli for mentoring and supporting diatom identification. We thank Tania Rivlin for inorganic nutrient measurements and Eyal Rahav, Wiebe Kooistra and Chana Kranzler for feedback comments on the study.

## Data Archiving

Data is available from the authors upon request.

