## Supplementary material for "Diatom community shifts across oligotrophic-eutrophic seasonal gradients and bloom variability as a function of mixing depth in the subtropical Northern Red Sea": Supp. Fig. 1

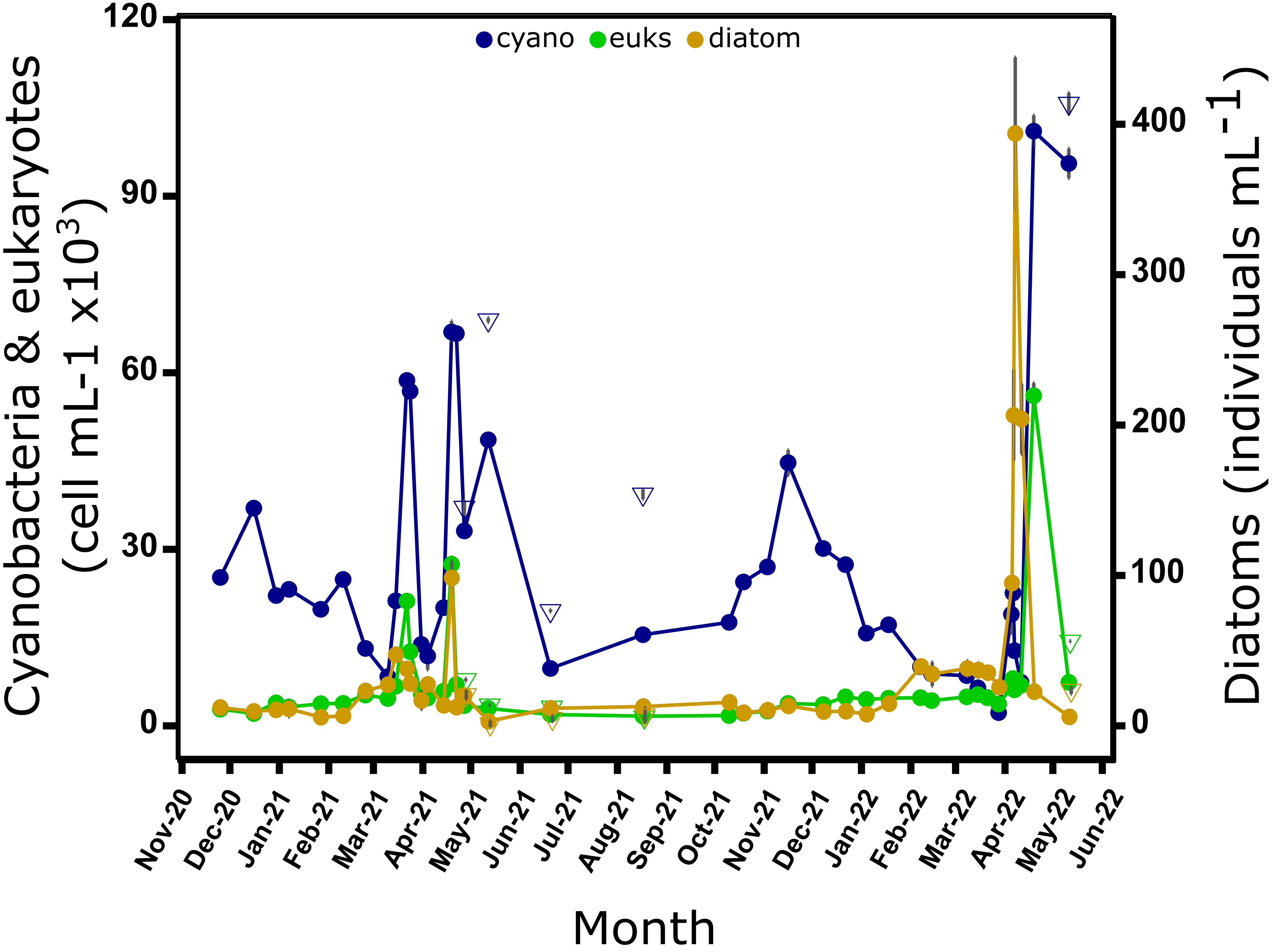


**Supplementary Figure 1.** Averaged concentration of total eukaryotic phytoplankton, total cyanobacteria (*Prochlorococcus* and *Synechococcus*) and diatom densities (n = 2). Circles connected with line represent 2m depth, while triangles represent samples from the deep chlorophyll maximum (DCM). Data of cyanobacteria and eukaryotes is based on flow cytometry counts, diatoms were counted by scanning electron microscopy.
