## Supplementary material for "Diatom community shifts across oligotrophic-eutrophic seasonal gradients and bloom variability as a function of mixing depth in the subtropical Northern Red Sea": Supp. Fig. 2

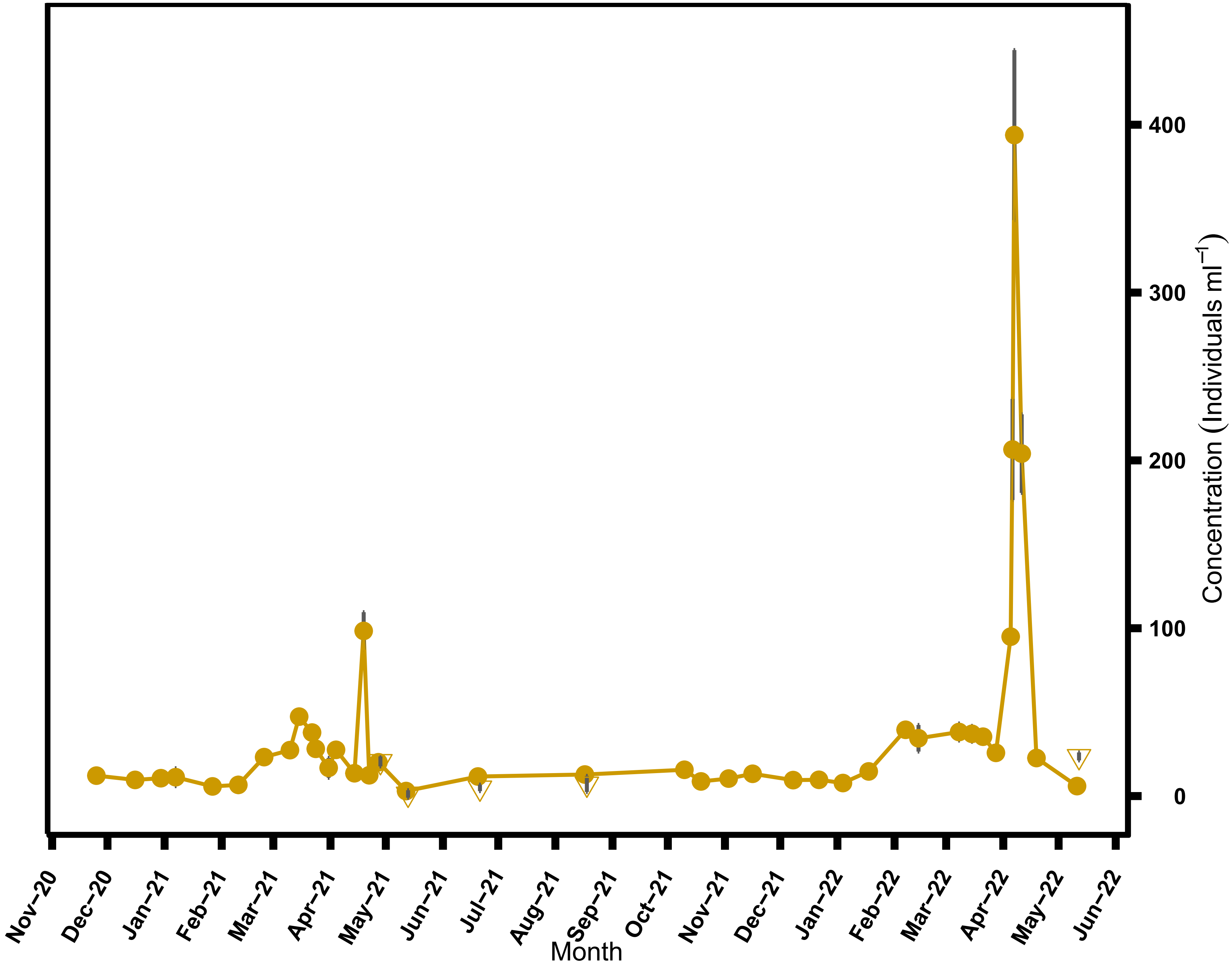


**Supplementary Figure 2.** Diatom abundance by SEM counts. Circles connected with line represent 2m depth, while triangles represent samples from the deep chlorophyll maximum (DCM). Grey bars represent standard deviation of diatom counts (n=2).
