## Supplementary material for "Diatom community shifts across oligotrophic-eutrophic seasonal gradients and bloom variability as a function of mixing depth in the subtropical Northern Red Sea": Supp. Fig. 3

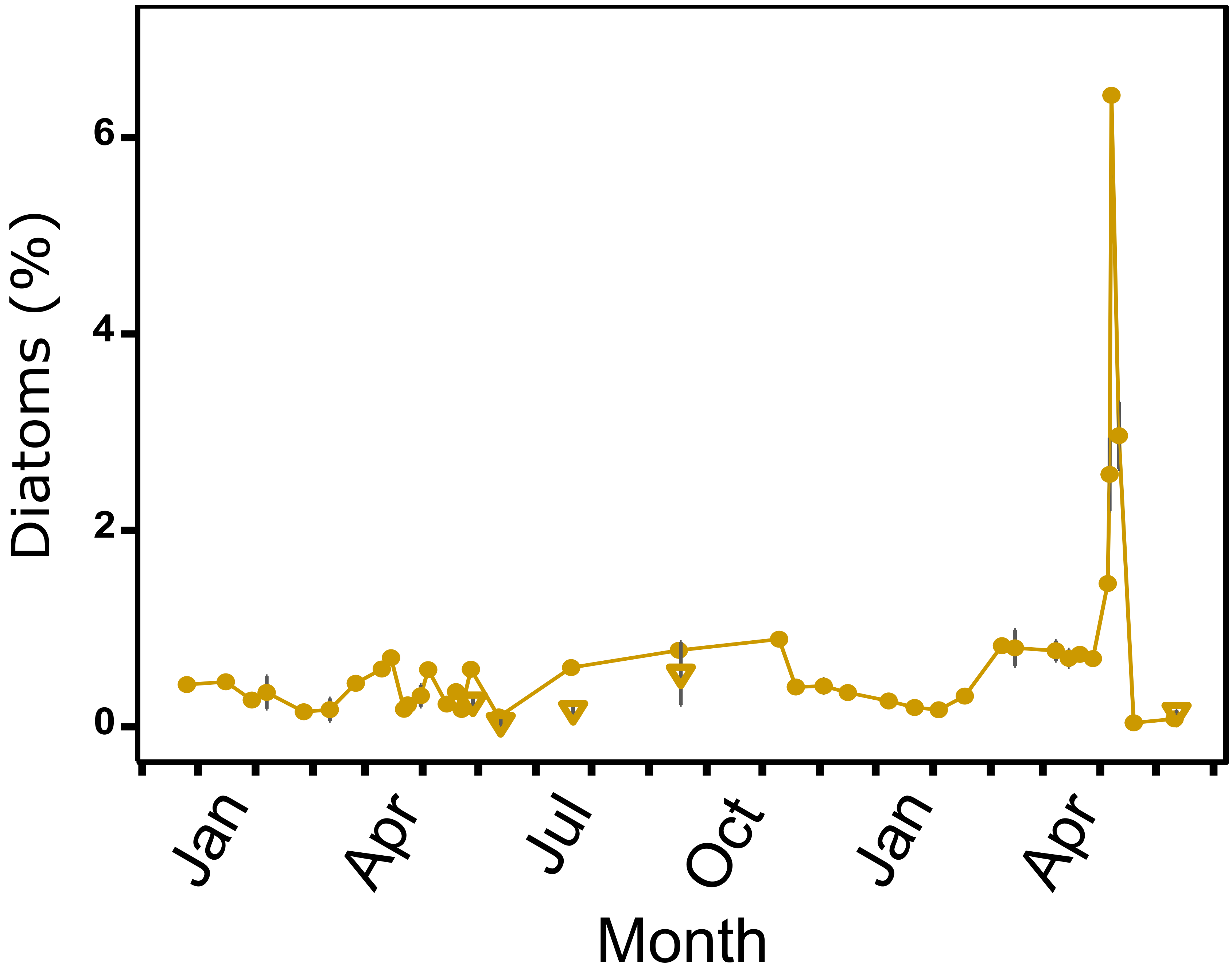


**Supplementary Figure 3.** Percentage of diatom counts (SEM) from total eukaryotes (flow cytometer). Circles connected with line represent 2m depth, while triangles represent samples from the deep chlorophyll maximum (DCM). Grey bars represent standard deviation of diatom counts (n=2), compared to average eukaryotes concentration (n=3).
