## Supplementary material for "Diatom community shifts across oligotrophic-eutrophic seasonal gradients and bloom variability as a function of mixing depth in the subtropical Northern Red Sea": Supp. Fig. 4

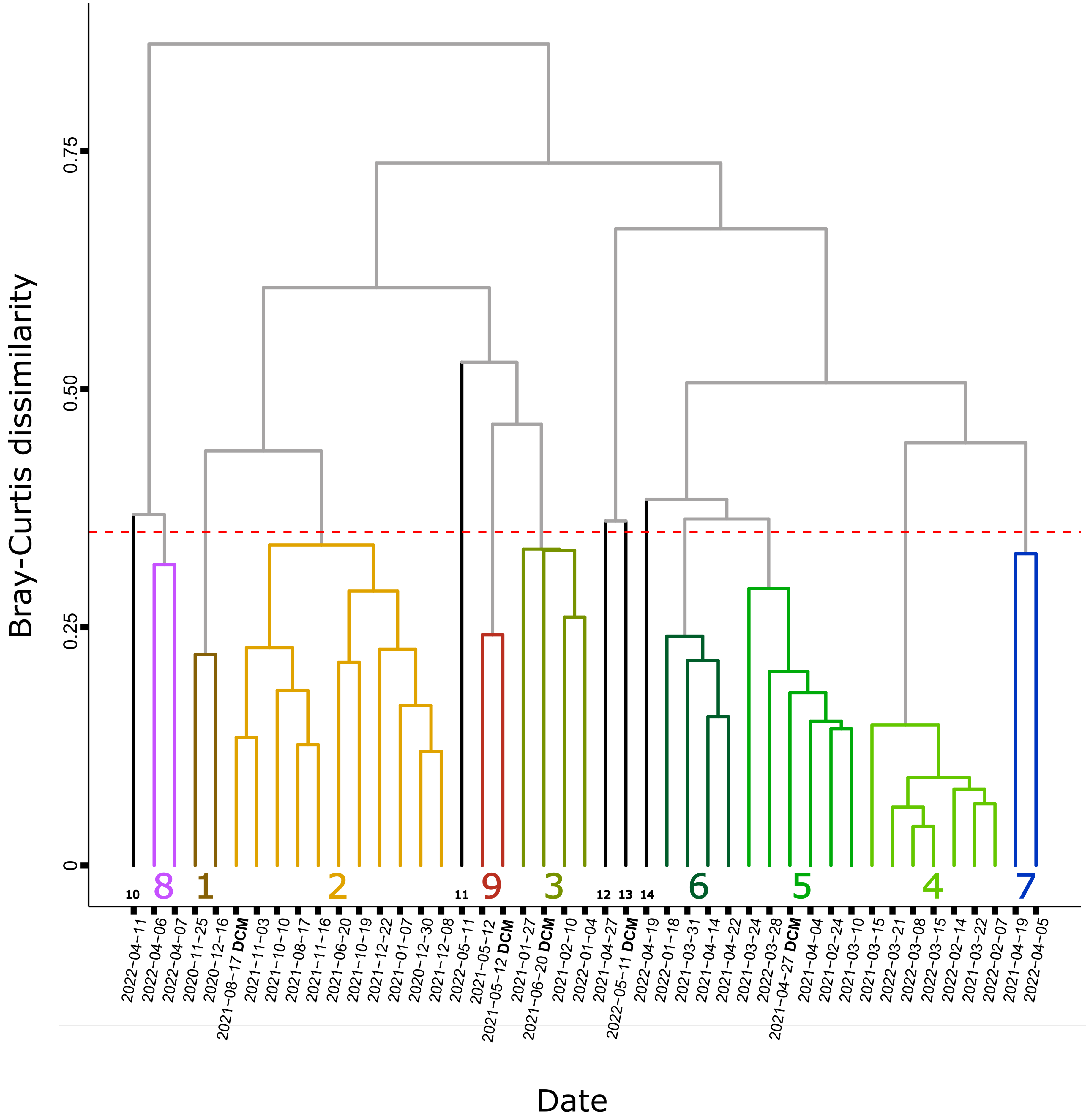


**Supplementary Figure 4.** Cluster analysis of diatom communities, based on Bray-Curtis dissimilarity. Red dashed line indicates 65% similarity, which served as cut-off for clustering. Branches are colored according to the 9 resulting communities, which served as the base for NMDS ordination. Numbers represent each community: 1,2: Summer-Autumn. 3-5: Mixing. 6: Spring transition. 7-8: Bloom. 9: Bloom crash. Black branches represent communities with low similarity (< 65%).
