## Supplementary material for "Diatom community shifts across oligotrophic-eutrophic seasonal gradients and bloom variability as a function of mixing depth in the subtropical Northern Red Sea": Supp. Table 1

Supplementary Table 1. Main species in each cluster

| Cluster | species | avg. C (Individual mL$^{-1}$) |
|---|---|---|
| 1 | *Cylindrotheca* spp. | 3.2 |
| | Thalassionemataceae | 1.9 |
| | *Pseudo-nitzschia* spp. | 1.7 |
| | 20>Pennate | 1.3 |
| | 50>Pennate>20 | 1 |
| 2 | *Cylindrotheca* spp. | 6.6 |
| | 50>Pennate>20 | 1.4 |
| | 20>Pennate | 1.3 |
| | *Pseudo-nitzschia* spp. | 0.4 |
| | Thalassiosiraceae | 0.4 |
| 3 | Thalassiosiraceae | 1.8 |
| | 20>Pennate | 1.3 |
| | *Cylindrotheca* spp. | 1.1 |
| | 50>Pennate>20 | 0.5 |
| | *N. hasleae* | 0.4 |
| 4 | Thalassiosiraceae | 16 |
| | *Cylindrotheca* spp. | 3.4 |
| | 20>Pennate | 2.6 |
| | *Pseudo-nitzschia* spp. | 1 |
| | 50>Pennate>20 | 0.9 |
| 5 | Thalassiosiraceae | 32.9 |
| | 20>Pennate | 1.6 |
| | *Cylindrotheca* spp. | 1.2 |
| | *Pseudo-nitzschia* spp. | 0.6 |
| | 50>Pennate>20 | 0.5 |
| 6 | Thalassiosiraceae | 7.7 |
| | *Cylindrotheca* spp. | 2.2 |
| | 20>Pennate | 2.1 |
| | 50>Pennate>20 | 0.9 |
| | *Pseudo-nitzschia* spp. | 0.6 |
| 7 | Thalassiosiraceae | 51.7 |
| | *Pseudo-nitzschia* spp. | 14.9 |
| | *Chaetoceros* spp. | 10.9 |
| | 20>Pennate | 8.5 |
| | 50>Pennate>20 | 5.1 |
| 8 | 20>Pennate | 0.8 |
| | *Cylindrotheca* spp. | 0.8 |
| | Thalassiosiraceae | 0.4 |
| | *Pseudo-nitzschia* spp. | 0.3 |
| | 50>Pennate>20 | 0.1 |
| | Thalassiosiraceae | 95.9 |
| | *Pseudo-nitzschia* spp. | 72.8 |

| | | |
|---|---|---|
| 9 | *Leptocylindrus sp.* | 49.7 |
| | *Chaetoceros* spp. | 44.9 |
| | 50>Pennate>20 | 13 |
| Un-clustered samples | | |
| 10 | 50>Pennate>20 | 6 |
| | 20>Pennate | 4.3 |
| | Thalassiosiraceae | 3.9 |
| | *Pseudo-nitzschia* spp. | 2.9 |
| | *Chaetoceros* spp. | 1.1 |
| 11 | Thalassiosiraceae | 123.3 |
| | *Leptocylindrus sp.* | 23.6 |
| | *Pseudo-nitzschia* spp. | 16.3 |
| | 50>Pennate>20 | 14.4 |
| | *Chaetoceros* spp. | 6.2 |
| 12 | Thalassiosiraceae | 10.4 |
| | Centric ungrouped | 5.5 |
| | 50>Pennate>20 | 2.7 |
| | 20>Pennate | 1.9 |
| | *Pseudo-nitzschia* spp. | 1.6 |
| 13 | 20>Pennate | 2.8 |
| | 50>Pennate>20 | 1.2 |
| | *Cylindrotheca* spp. | 0.9 |
| | Centric ungrouped | 0.8 |
| | Thalassiosiraceae | 0.2 |
| 14 | 20>Pennate | 7.9 |
| | 50>Pennate>20 | 6.8 |
| | Centric ungrouped | 6 |
| | Thalassiosiraceae | 2.3 |
| | *Cylindrotheca* spp. | 1.2 |
